# A Homology Independent Sequence Replacement Strategy in Human Cells Using a CRISPR Nuclease

**DOI:** 10.1101/2020.05.11.088252

**Authors:** Eric Danner, Mikhail Lebedin, Kathrin de la Rosa, Ralf Kühn

**Author notes:** Corresponding Authors ED, RK.

## Abstract

Precision genomic alterations largely rely on Homology Directed Repair (HDR), but targeting without homology using the Non-Homologous End Joining (NHEJ) pathway has gained attention as a promising alternative. Previous studies demonstrated precise insertions formed by the ligation of donor DNA into a targeted genomic double strand break in both dividing and non-dividing cells. Here we extend this idea and use NHEJ repair to replace genomic segments with donor sequences; we name this method ‘Replace’ editing (<u>R</u>ational <u>e</u>nd-joining <u>p</u>rotocol de<u>l</u>ivering <u>a</u> targeted sequen<u>c</u>e <u>e</u>xchange). Using CRISPR/Cas9 we create two genomic breaks and ligate a donor sequence in-between. This exchange of a genomic for a donor sequence uses neither microhomology nor homology arms. We target four loci and show successful exchange of exons in 16% to 54% of cells. Using linear amplification methods and deep sequencing pipelines we quantify the diversity of outcomes following Replace editing and profile mutations formed at the ligated interfaces. The ability to replace exons or other genomic sequences in cells not efficiently modified by HDR holds promise for both basic research and medicine.

## Introduction

RNA guided nucleases (1–3) have rapidly become foundational tools in facilitating genomic manipulations (4, 5). These nucleases target specific genomic loci and form a double strand break (DSB). DNA repair processes are then leveraged to produce the desired outcome of the gene editing. Conventionally, specific genomic changes are made using Homology Directed Repair (HDR) (6, 7) with exogenously introduced DNA containing flanking sequences homologous to the targeted locus. One limitation of HDR mediated genome editing is its restriction to the S/G2 phase, reducing or abolishing efficacy in slowly or non-dividing cells (8). Additionally, HDR can be precise but recent reports demonstrate greater error than often assumed, as incomplete or extraneous portions of the delivery vector are copied into the genome (9, 10). On the other hand, the canonical Non-Homologous End Joining (NHEJ) pathway is traditionally viewed as error prone and relegated to disrupting gene function by inducing small insertions and deletions (InDels) during DSB repair. However, the high fidelity aspects of NHEJ repair are often underappreciated as mutant InDels are easily observed, whereas non-mutagenic repair is indistinguishable from the original allele (11). Furthermore non-mutagenic repair by NHEJ reforms the Cas9 target site allowing for continued DSB formation. This may result in a final genomic population containing majority InDels despite NHEJ repair being predominately error free.

Recently, increasing awareness of the fidelity and efficiency of NHEJ repair has led to the development of methods to produce genomic deletions and exogenous sequence insertions using this pathway. As NHEJ is highly active in all phases of the cell cycle, it has been used for gene editing in dividing and non-dividing cells, such as muscles and neurons (12–14). Targeted deletions are produced by forming two DSBs with loss of the intervening sequence during repair. The ubiquitous nature of the NHEJ pathway allows for deletions in zygotes, as well as in adult tissue such as *in vivo* exon deletion in a mouse muscular dystrophy model (14, 15). Additionally, exogenously introduced dsDNA donor sequences can efficiently ligate into a single DSB by NHEJ (herein referred to as Insert targeting) (12, 13, 16–23). With the NHEJ pathway conserved broadly, Insert targeting has been shown in plants (23), fish (18), cell lines (16, 17, 19–22), human iPSC-derived neurons, and *in vivo* mouse tissues (12, 13). The ability to effectively integrate DNA across cell types has been used to tag genes with fluorophores (12, 16, 22), identify off-target CRISPR cleavage sites (24), and as a strategy for gene therapy by inserting functional coding sequences upstream of a disease causing exon (13).

The ability to leverage NHEJ repair to create large deletions and insert exogenous DNA posits the possibility of NHEJ-based sequence replacement; two DSBs are produced and a donor sequence without homology is ligated between the two breaks. This approach would enable the replacement of defective exons or regulatory sequences in a wide range of resting or dividing cells. While NHEJ-based replacement has been demonstrated in plants, where HDR is often infeasible (25, 26), it has not yet been applied in animal cells. Here we demonstrate efficient replacement of genomic sequences with a donor sequence in human cells using NHEJ repair; we call this method Replace (<u>R</u>ational <u>e</u>ndjoining <u>p</u>rotocol delivering <u>a</u> targeted sequen<u>c</u>e <u>e</u>xchange). Using fluorescence models we demonstrate efficient Replace editing and the exchange of exons in multiple genes. The structural variants produced during Replace targeting motivated the development of sequencing pipelines to better quantify the results. This demonstration provides a proof of concept for NHEJ-based sequence replacement in human cells and provides design principles to guide future applications in gene therapy and research.

## Results

For initial testing of Replace targeting we used a fluorescence based reporter system. The synthetic reporter system was created and integrated into the AAVS1 locus in HeLa cells (Fig. 1A). The reporter system contains a CAG promoter upstream of a BFP fluorophore. A polyadenylation sequence (pA) following the BFP prevents the expression of a downstream Venus-pA. The cells initially are BFP^+^. Replace targeting exchanges the BFP cassette with a mCherry donor. Reporter HeLa cells were lipofected with the donor and a Cas9 plasmid containing puromycin resistance gene. Lipofected cells were puromycin selected for 48 hr and cells were analyzed two weeks later (Fig. 1B). Replacement was caused by SpyCas9 cleavage on both sides of the BFP-pA cassette which was then free to exchange with the linearized mChery donor sequence. Correct ligation of mCherry resulted in mCherry^+^ cells. Excision of the BFP cassette without replacement, a deletion, resulted in expression of the downstream Venus-pA. Some alleles lost expression due to deleterious resection or incorrect donor ligation. For comparison, Insert targeting and HDR were also carried out in an identical setup using a single gRNA targeting upstream of the BFP and a donor sequence with or without homology arms, respectively (Fig. 1A).

**Figure 1:**
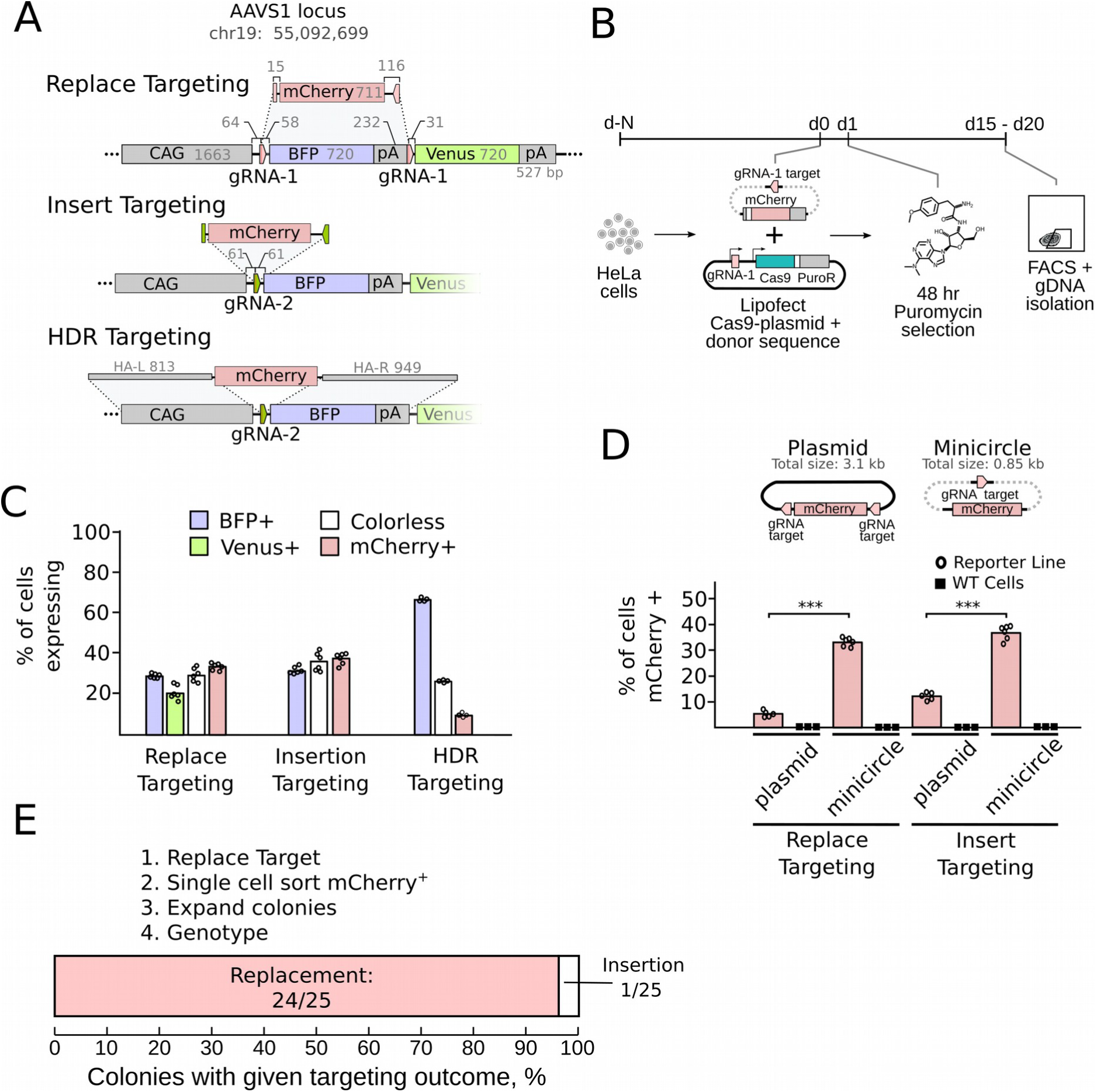
Initial validation of Replace targeting using HeLa reporter cells. **A.** HeLa cells with two copies of the fluorescent reporter system integrated into AAVS1 can be modified with Replace, Insert, or HDR targeting. Gray numbers are sequence lengths in base pairs (bp). **B.** Experimental scheme showing Replace targeting. Lipofection of Cas9-2A-puro/gRNA-1 plasmid and mCherry donor sequence into HeLa reporter cells followed by 48 h puromycin selection. All Replace, Insert, HDR targeting followed the same setup but with corresponding plasmids. **C**. Results of targeting as measured by FACS of transfected reporter cells. Replace and Insert targeting were performed using minicircle donor sequences. Totals correspond to more than 100% as cells have two loci and may express two fluorophores. **D.** Replace targeting using minicircles and plasmid sequence donors. Replace targeting uses gRNA-1 sites to linearize plasmids or minicircles, and cuts the genomic site. Insert targeting correspondingly uses gRNA-2 sites. WT cells are HeLa cells without the reporter system in AAVS1. They are targeted identically to the reporter line. P-value were calculated using Student’s t-test. (**** P-value < 0.0001). **E.** Replace targeted, mCherry^+^ cells were single cell sorted, expanded, and genotyped. Correct replacement of BFP by mCherry (Replacement) and mCherry integration flanking the BFP (Insert) were both measured.

Replace targeting of the reporter locus resulted in 34% mCherry^+^ cells (Fig. 1C). Similarly, NHEJ-mediated Insert targeting resulted in 37%, mCherry^+^ cells, whereas HDR produced only 9% mCherry^+^ cells and most cells remained BFP^+^ (67%). We compared the effect of delivering donor sequences within a plasmid or in the form of minicircles as a previous report showed minicircles to increase Insert efficiency (13) (Fig. 1D). Minicircles are minimal plasmids and contain only the donor sequence. Thus they require only a single Cas9 DSB for linearization, while plasmid delivery requires two Cas9 cuts to excise the donor. Donor sequences delivered as minicircles resulted in a six-fold increase in Replace targeting compared to plasmid delivery. We therefore used minicircles for Replace targeting in the remainder of this work. To address if mCherry expression was driven in part by off-target integration of the donor sequence, we Replace targeted, in an otherwise identical manner, wild-type HeLa cells. As these cells do not contain the AAVS1 integrated promoter and target site, only off-target integration could result in mCherry expression (Fig. 1D). Wild type HeLa cells showed no mCherry expression indicating that the 34% mCherry^+^ cells in our original experiment are the result of integration at the target loci of our reporter system. mCherry^+^ cells were single-cell sorted, expanded, and genotyped to check for correct sequence replacement. 24 out of 25 analyzed clones, i.e. 32% of all cells, contained the anticipated replacement of BFP with mCherry, while one clone contained an allele with mCherry insertion upstream of BFP (Fig. 1E). As HeLa reporter cells contained two copies of the reporter locus we worked to quantify the frequency of homozygous knock-in. To estimate this population we simultaneously transfected two donor sequences (mCherry and miRFP670) (Suppl. Fig. 1A,1B). By measuring the mCherry^+^, RFP^+^, and dual positive populations we calculated an average of 5% homozygous knock-in (Suppl. Fig. 1C, 1D). Taken together, Replace targeting in our reporter system occurs as a major outcome, with a successful sequence exchange in 32% of cells.

During the ligation of the donor sequence into the genome, InDels may occur at the genome-donor sequence interface. To ascertain the major InDel outcomes, the gDNA of targeted, unsorted HeLa reporter cells was PCR amplified using primers flanking the ligated interface. The deconvolution of the Sanger traces of these amplicons provides an InDel estimate of the bulk population of Replace (Fig 2A) and Insert targeted (Fig. 2B) alleles. Overall this analysis showed that resection occurred as a minor subset (<16%) of the ligated interfaces in Replace or Insert targeted alleles. The major product was either ligation of the donor into the genome without producing an InDel, or small (1-2 nucleotide) insertions. Sanger sequencing of cloned individual alleles supports the bulk analysis (Suppl. Fig. 2). The one or two nucleotide insertions were striking in that they were non-random and matched the protospacer sequence downstream of the break site. It is known that SpyCas9 does not always form its canonical blunt end break 3 nucleotides downstream of the PAM, but can, at some frequency, form a staggered cut (27–30). We believe that the formation of the non-random insertion InDels is caused by this non-canonical sticky-end cutting of SpyCas9 and not by NHEJ (Fig. 2C). In this model sticky end cutting causes the PAM side (PAMside) of the break to contain nucleotides normally present on the protospacer side (protoside). These overhangs are filled during repair and appear as insertions when the PAMsides are ligated. The design of Replace and Insert sequence donor results in two PAMsides or two protosides being ligated in the final product (Suppl. Fig. 3). These insertions then are therefor seen only on the PAMside-PAMside interface.

**Figure 2:**
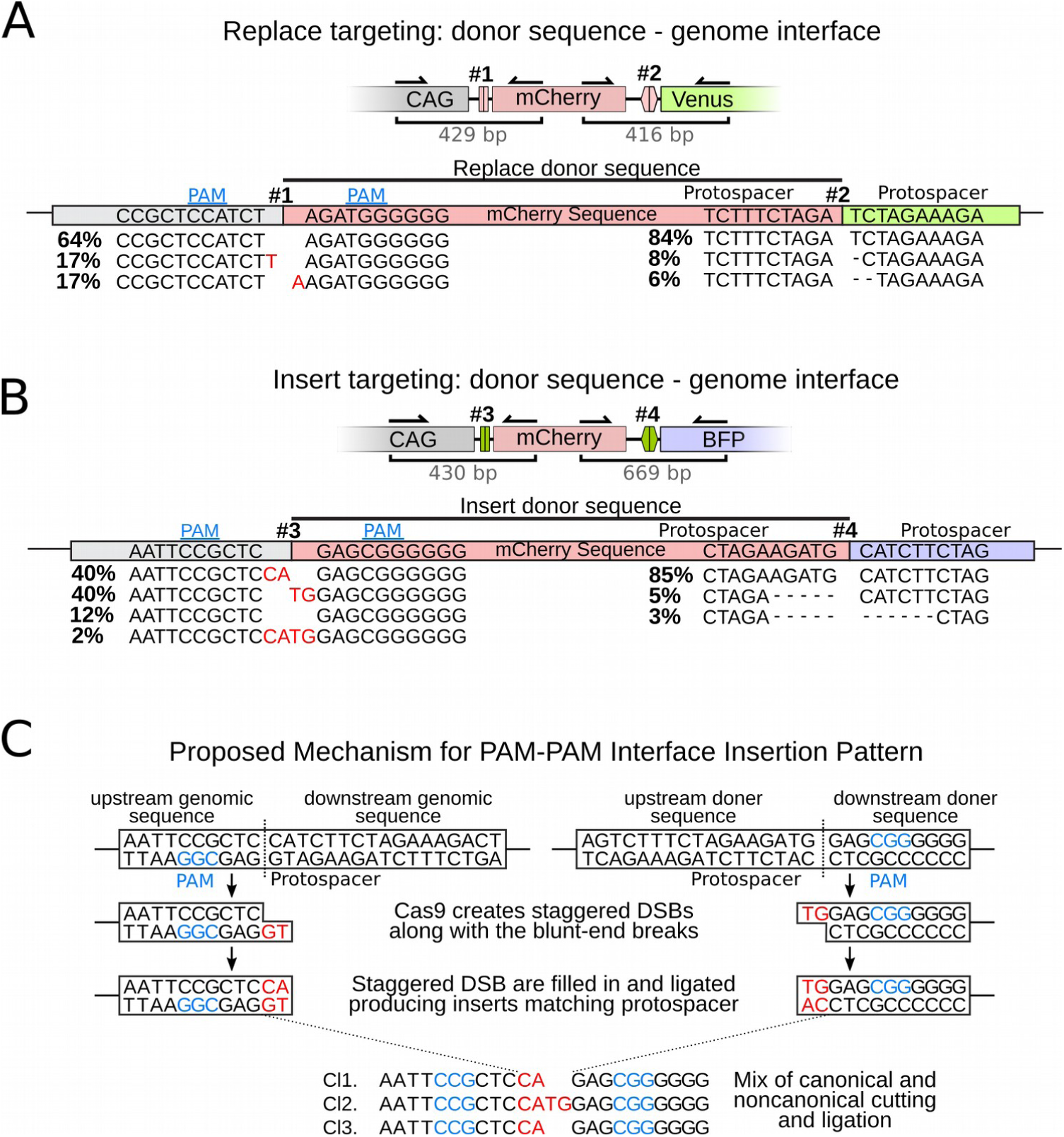
Sequencing for InDels at donor sequence-genome junction in edited HeLa reporter alleles. **A.** InDel frequencies at the interface of the integrated donor sequence in Replace targeted cells quantified by ICE deconvolution. **B.** InDel frequencies at the interface of integrated donor sequences following Insert targeting quantified by ICE deconvolution. Recurrent insertion pattern highlighted in red. **C.** Proposed mechanism driving recurrent insertion pattern at PAM-PAM junctions using HeLa Reporter Insert targeting data.

It is known that large scale resection may follow a single Cas9 driven DSB (31), and Replace targeting further complicates analysis due to the structural variants formed by the two genomic breaks and donor sequence integration. In order to quantify resection and the directionality of donor integration, we performed long-read deep-sequencing on amplicons of the Replace targeted loci of unsorted HeLa reporter cells using primers 800-900 bp away from the DSB sites. A bioinformatics pipeline was built to analyze resection and structural outcomes frequencies (Fig. 3A) (Suppl. Fig. 4 A,B). Alignment of the reads showed alleles with >500 bp of resection occurred (Fig. 3B). Notably, individual reads showed that large scale resection was frequently asymmetric with one side of the break undergoing dramatically larger resection. Viewing the average resection frequency at each base along the amplicon showed that in the majority of cases resection was modest (Fig. 3C). Specifically, in alleles with successful mCherry replacement of BFP, the resection at the ligated junctions was smaller than 10 bp in 85% of the reads, and Protoside-Protoside junctions were completely InDel-free in 66% of the reads. The donor sequence has no homology and is expected to integrate equally in both directions. However, inspired by from the work of Suzuki. et. al. (13), we designed a preferred orientation into our donor sequence (Suppl. Fig. 5). Constructs are designed so that donor sequences, when integrated in the undesired direction, form a Cas9 target site, whereas donor integration in the desired orientation abolished further Cas9 cutting. Analysis of the long read data show that alleles with successful BFP replacement integrated mCherry in the designed orientation in 79% of the cases (Fig. 3D). Even alleles containing unintended donor insertion onto either side of the BFP contained the mCherry in the designed orientation in 67% of the cases.

**Figure 3:**
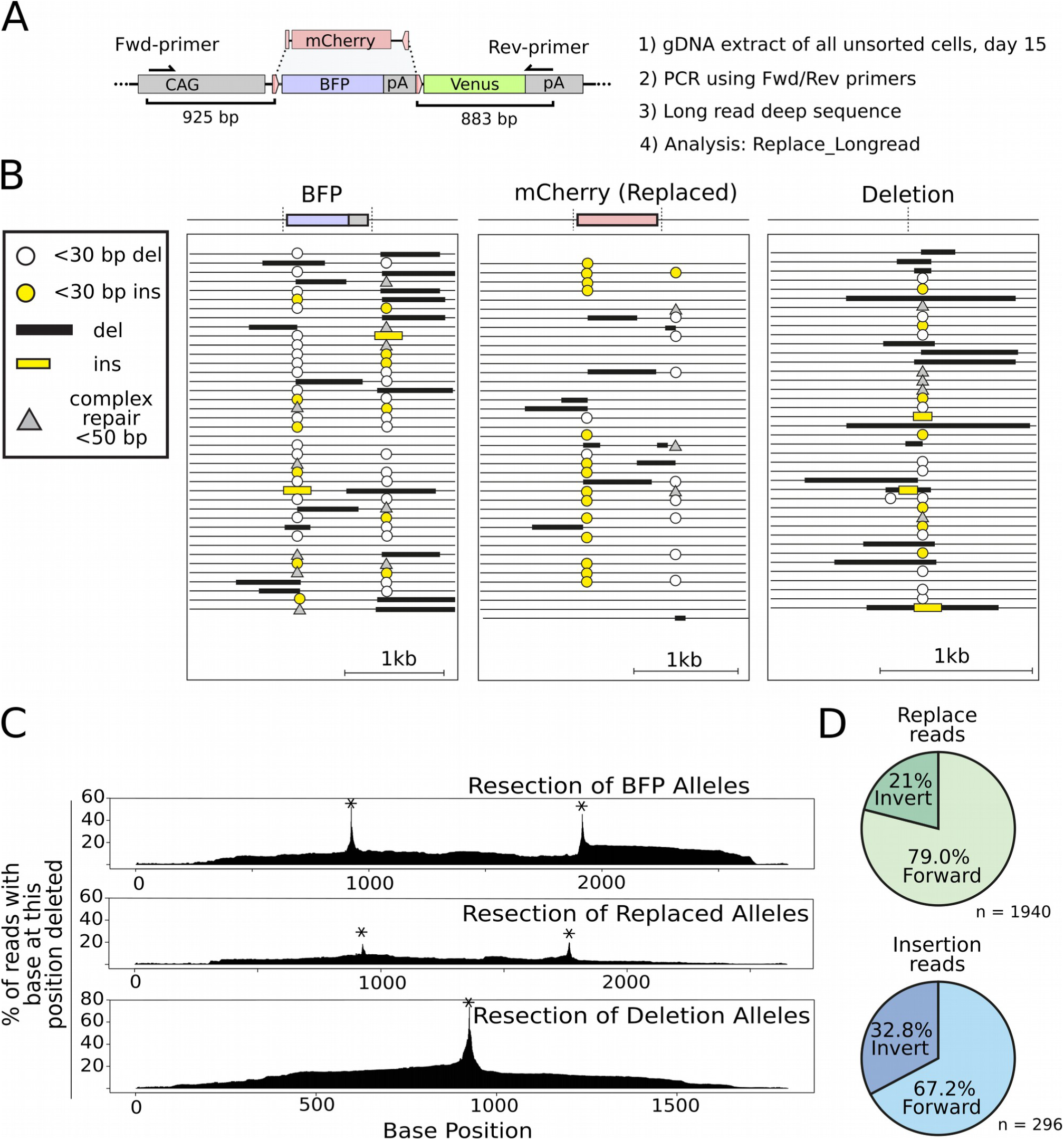
Long read deep sequencing of Replace targeted HeLa reporter cells. **A.** Unsorted cells after Replace targeting were used for gDNA extraction, loci amplification, PacBio sequencing, and sequence analysis. **B.** Forty representative alignments of three major type of alleles: the original BFP exon, the replacement with mCherry, or BFP deletion. **C.** Resection profile of alignments containing only the original BFP sequence, the donor mCherry sequence (Replaced), or BFP deletions. Alignment files were analyzed for the fraction of reads containing a deletion at each base for a given allele set. The plot represents an average resection profile of the alignment. Asterisk denotes Cas9 cleavage site. **D.** Directionality of structural variants formed after Replace targeting.

To test Replace targeting of an exon from a natural gene, we targeted the ubiquitously expressed Polymerase Beta (*POLB)* gene in K562 cells. We replaced exon 5 with a splice acceptor-2A-mCherry-pA donor sequence (Fig. 4A, 4B). Replace targeting resulted in 58% of cells being mCherry^+^ on average, with reporter expression stable over weeks (Fig. 4C, 4D). Genotyping of mCherry^+^ single-cell derived colonies showed mCherry integration into the *POLB* locus in 100% of colonies and correct replacement of exon 5 in 51/55 cells, with 4/55 containing both the mCherry donor sequence and the original exon (Fig. 4E). Sanger trace deconvolution of the genome-donor sequence interface and individual PCR clone reads showed modest InDel formation in the replaced alleles (Fig. 4F)(Suppl. Fig. 2). Furthermore, *CCNA1* exon 2 and *LMNA* exon 2 were also Replace targeted using the same mCherry donor sequence backbone. These Replace targeting tests resulted in 39% and 19% mCherry^+^ cells respectively (Fig. 4G, 4H). Genotyping of mCherry^+^ single cell colonies showed replacement as the dominant outcome, occurring in 60% of *CCNA1* colonies and 85% of *LMNA* colonies analyzed (Suppl. Fig. 6, Suppl. Fig. 7). Combining FACS and single cell genotyping data allowed an estimate of 54% of *POLB*, 23% of *CCNA1*, and 16% of *LMNA* Replace targeted cells with successful replacement.

**Figure 4:**
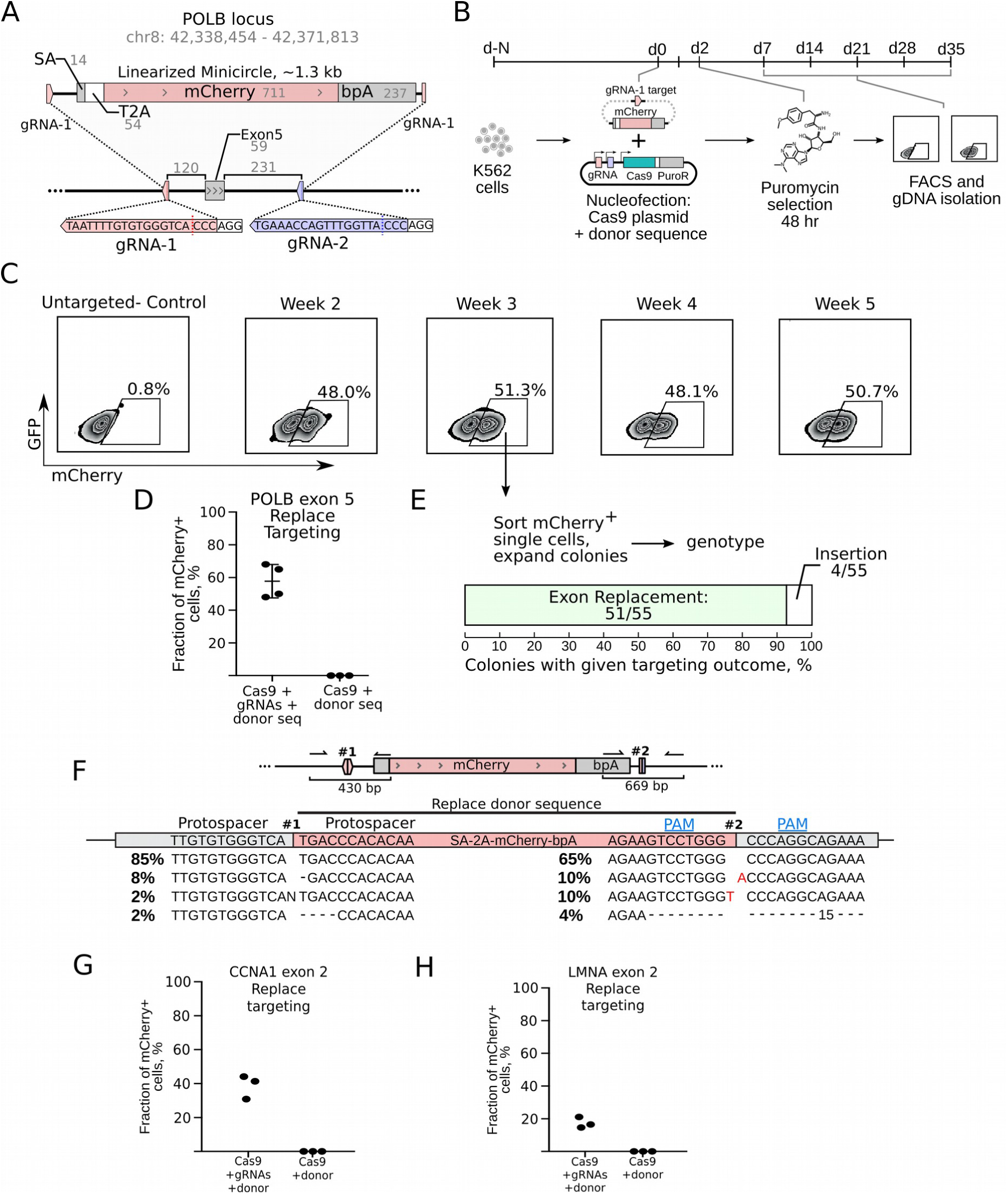
Exon Replace editing in K562 cells. **A.** Replacement of *POLB* Exon 5 with a cherry reporter including a splice acceptor (SA), T2A self-cleaving peptide and a polyadenylation site (bpA). Gray numbers represent sequence length in base pairs (bp). **B.** K562 cells cotransfected with Cas9-2A-puro/guides plasmid and mCherry Minicircles were puromycin selected for 48h and followed by FACS analysis over 5 weeks. **C.** mCherry^+^ expression in transfected cell cultures was measured weekly by FACS. **D.** Biological replicates of Replace targeting of *POLB* Exon 5. **E.** Colonies of mCherry^+^ single cells were expanded and genotyped to check and quantify replacement of the exon with the donor sequence. **F.** InDel frequencies at the interface of the integrated donor sequence in Replace targeted unsorted cells quantified by Sanger trance deconvolution of amplicons #1 or #2. **G.** Replace targeting *CCNA1* exon 2 in K562 cells with SA-2A-mCherry-pA donor. **H.** Replace targeting of *LMNA* exon 2 with a SA-2A-mCherry-pA donor.

To measure resection during exon exchange and to measure the directionality of integrated donor sequences, we long-read sequenced the *POLB* exon 5 targeted locus in the unsorted cells (Fig. 5A). The primers were 1-2kb away from the break site to better capture possible larger scale resection. The mCherry donor sequence was again designed to integrate in a specific orientation (Suppl. Fig. 5). Analysis of all measured alleles showed 89% of alleles with exon 5 replaced by mCherry were in the designed orientation (Fig. 5B). Donor sequences even integrated preferentially in alleles containing the donor sequence inserted flanking exon 5. Resection was closely examined as resection could damage the splicing sequence or coding sequence of the donor sequence. Large InDels at the ligated interfaces were infrequent with >95% of the correctly replaced exon 5 reads containing less than 30 bases of resection at either end. Strikingly, nearly half of the correctly targeted replacement reads had no InDels at the Protoside-Protoside interface (Fig. 5C, 5D).

**Figure 5:**
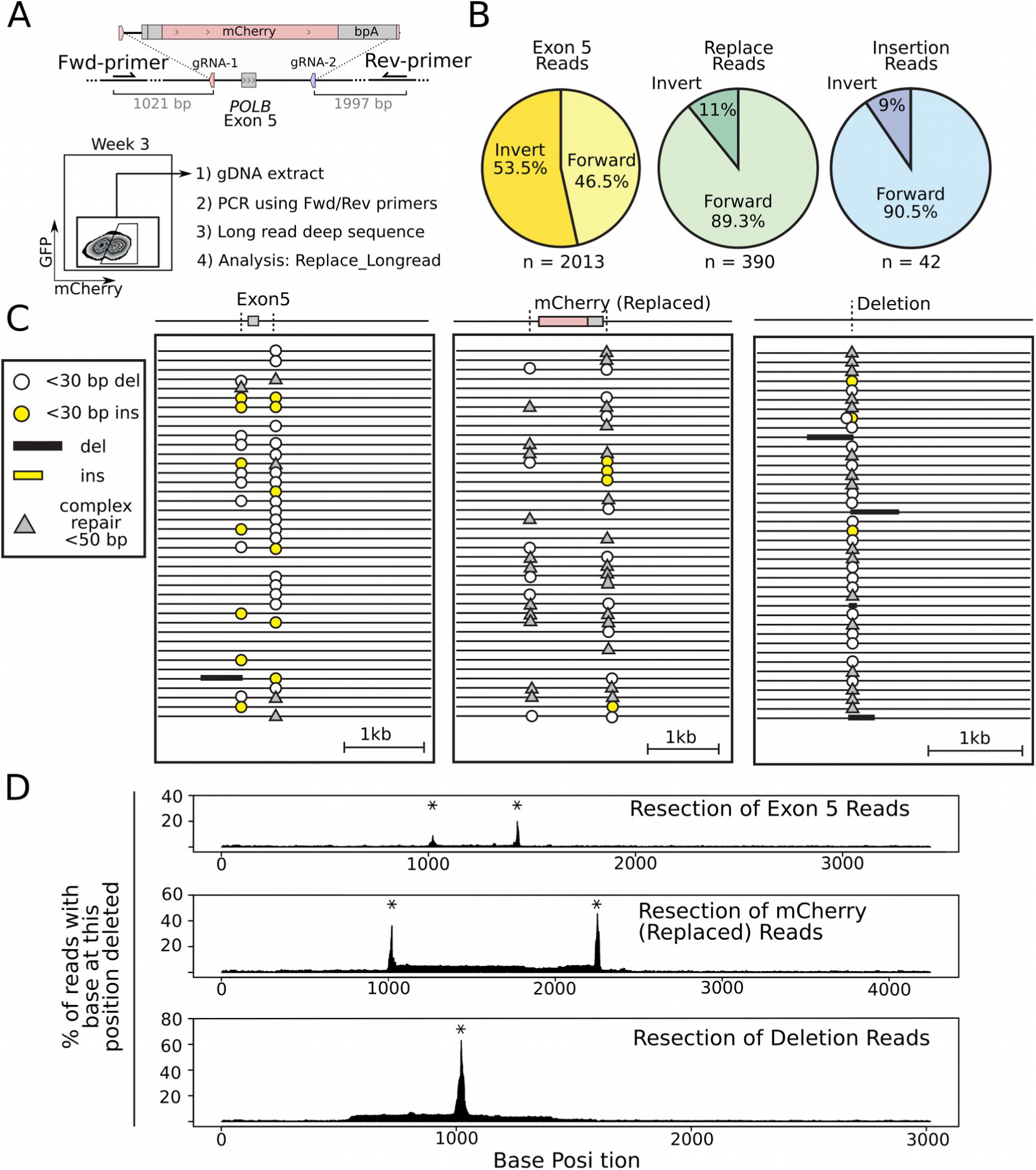
Long read deep sequencing analysis of *POLB* exon 5 after Replace editing in K562 cells. **A.** Long read deep sequencing of unsorted *POLB* exon 5 Replace targeted K562 cells. Loci were amplified using the primers more than 1 kb from the Cas9 breaksites; amplicons were sequenced using PacBio technology. **B.** Directionality of structural variants following Replace targeting. **C.** Forty representative alignments of three major type of alleles: exon 5 retaining allele, exon replacement with mCherry, or a deletion allele. **D.** Resection profile of alignments containing the original exon 5 sequence, the donor mCherry (Replaced) sequence, or deletions. Plots represent average resection profiles of the measured alleles. The percentage of alleles containing a deletion at each individual base position were calculated from the alignments in a given allele set with asterisk marking the location of Cas9 target site.

Linear PCR methods requiring only one gene specific primer, such as UdiTaS (32) and LAM-HTGTS (33), offer more complete and quantitative measurements of DNA repair outcomes following a DSB. A gene specific primer binds upstream of the targeted break site and a universal primer binding sequence is integrated downstream. Subsequently, the PCR amplifies the region across the break regardless of the structural variant, deletion size, or translocation (Fig. 6A) (Suppl. Fig. 4C, 4D). The UDiTaS method also contains a robust computational pipeline for CRISPR analysis. We modified this pipeline to UDiTaS-Replace, extending the capabilities for Replace targeting with two pipelines (Suppl. Fig. 4E). Pipeline 1 closely follows the published UDiTaS pipeline; it aligns reads to the *in silico* reconstructed expected outcomes, performs InDel analysis, and quantifies these measurements. The results of Pipeline 1 showed that at the targeted *POLB* locus donor sequence integrated in the preferred orientation at a 5:1 ratio to an inverted orientation (Fig. 6A, 6B). At 39 % of all *POLB* alleles, the integration of the donor sequence in the desired orientation is the single most frequent outcome measured. Strikingly, more than ⅓ of these donors were integrated without an InDel formed at the ligated interface. This highlights both the efficiency and fidelity of Replace targeting for exon replacement.

**Figure 6:**
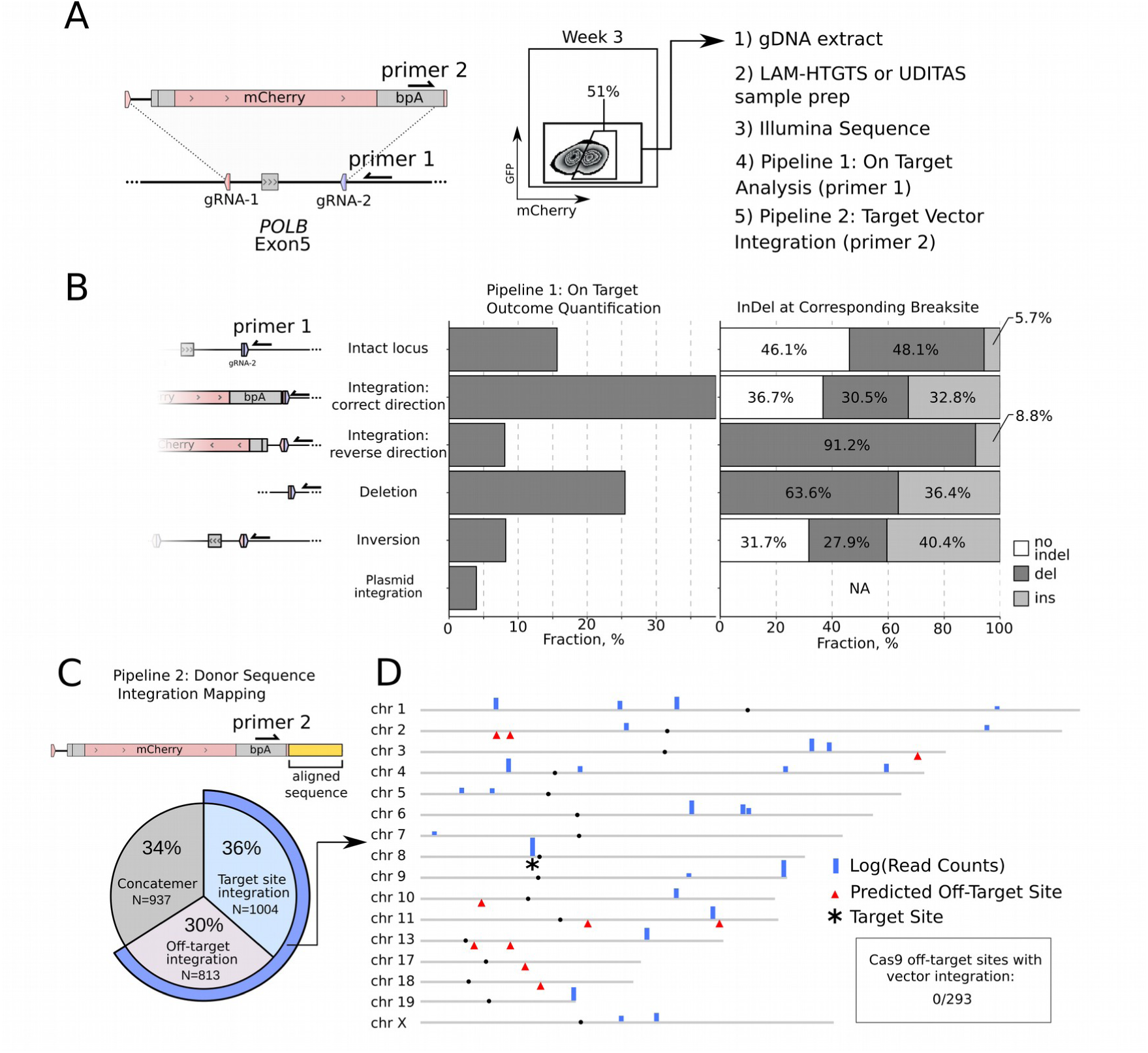
Linear amplification analysis of *POLB* exon 5; quantifying on-target Replace outcomes and mapping donor sequence integration. **A.** Replace overview showing primer binding used for linear amplification. Unsorted gDNA was amplified with primer 1 or primer 2 and a universal primer, sequenced using Illumina technology, and analyzed. **B.** Pipeline 1 analysis of Uditas prepared samples quantifies outcomes at the targeted site with the corresponding InDels quantification. **C.** Pipeline 2 analysis of Uditas prepared samples shows mCherry integration with an overall quantification of donor sequence integration. **D.** Donor sequence integration mapped across the genome with read counts plotted logarithmically. Chromosomes with no integration sites were removed. Top 10 predicted Cas9 off-target sites shown with red triangles. The *POLB* locus on chromosome (chr) 8 is marked by an asterisk.

As exogenously introduced DNA is known to integrate randomly into the genome (34), we developed UDiTaS-Replace Pipeline 2 to quantify and map the integration of the donor sequence (Suppl. Fig. 4F). Using a primer which binds to the donor sequence and points towards the ligation interface we generated amplicons that contain the flanking local genomic sequence (Fig. 6C). These amplicons were sequenced using Illumina technology, and the genomic sequences flanking the donor sequence break-site were aligned to the reference human genome (Fig. 6C, Suppl. Fig. 8, Suppl. Fig. 4C). Sequence alignment showed 55% on-target integrations into the *POLB* locus. 34% of all measured donor sequences had formed concatenations; it remains to be determined where these concatenated sequences are integrating within the genome. The donor sequences were shown to be integrated into the genome at more than 28 loci (Fig. 6D). Interestingly, none of the off-target integration mapped to any of the 293 predicted (35) SpyCas9 off-target sites.

## Discussion

This work is the first to demonstrate that NHEJ based genomic sequence exchanges are feasible and efficient in human cells. In the four loci tested replacement was successful in 16-54% of cells; in one case the desired product was the major outcome. We furthermore demonstrated targeted exon replacement via NHEJ in three widely expressed human genes. Based on the comprehensive analysis of our targeted alleles we arrive at three design principles to guide future Replace work.

The first design aspect ensures the correct orientation of the donor sequence in the genome. Linearizing the donor sequence with the same gRNA that cuts the target locus allows incorrectly ligated donors to be re-cut and excised. In addition, it is crucial to add a gRNA targeting the sequence formed during a deletion. This gRNA re-opens alleles that form a deletion and also excises out incorrectly ligated donor sequences (Suppl. Fig. 5). The minimal requirement for this design is two gRNAs (Suppl. Fig. 5B, 5C). Long read sequencing confirmed 89% of the donor sequences integrated in the designed orientation after *POLB* editing.

The second design principle is to avoid gRNAs that are involved in non-canonical SpyCas9 sticky end cutting. The frequency of ‘InDel free’ ligated interfaces measured in this work supports the idea that NHEJ repair is often not mutagenic (11). We believe breaks introduced by Cas9 are often re-ligated to reform the original sequence, which can then be cleaved again - forming a break ligation cycle. This cycle continues until the Cas9 is no longer active or the target site forms an InDel during repair and disrupts Cas9 binding. For efficient Replace or Insert targeting, prolongation of this cycle provides more time to acquire and ligate the donor sequence in the correct orientation. InDel mutations remove alleles from the ligation cycle and thus decrease efficiency. One avoidable driver of InDels formation is non-canonical SpyCas9 cutting in which a staggered cut is formed (27–30). The staggered cut is filled in and then ligated, duplicating the staggered nucleotide(s). The resulting small insertions are easily identifiable as they match the nucleotides of the protospacer sequence beyond the expected break site (Fig. 2C, Suppl. Fig. 3). Data from large gRNA screens suggest this mechanism as the predominant driver of +1 insertions (36). The non-canonical cutting of SpyCas9 may be sequence or loci dependent. Empirical testing of a gRNA by measuring InDel outcomes (37) therefore allows to avoid sites that incur staggered cuts. The benefits of blunt end cutting precludes RNA guided nucleases such as Cas12a that innately form staggered cuts. Cas9 nickases are subsequently also ill-suited for Replace targeting due to the staggered cuts formed and the high number of guides required.

The third concept is to design sacrificial sequences around the ligated regions to buffer possible resection. While the overall rate of InDels and resection is low, detrimental effects from resection can be further reduced. During exon Replace targeting we cut in intronic regions outside the splice site as short intronic InDels are less likely to be detrimental to gene function. Long range deep sequencing showed that in our systems the vast majority of the InDels are less than 30 bp long. Considering this, we recommend a sacrificial buffer 30 bp or greater be included on the flanks of the Replace construct to protect the splicing donor/acceptor and coding sequence. We currently use minicircles but also recommend such buffers on AAV delivered donor sequences too.

Measuring the outcomes of Replace targeting is complicated by the various structural rearrangements formed. Additionally, a growing body of literature documents complex outcomes following even simple Cas9 formed DSBs. These can include large scale resection (31), chromosomal fusions (32), mis-spliced mRNA, and unintended vector integration into the break site (14). In working towards a full understanding of the outcomes of Replace targeting we developed multiple deep sequencing pipelines. Long-read sequencing of PCR amplicons of the targeted loci proved useful in illuminating resection profiles and give insight into the orientation of the structural variants produced. However, samples prepared for long read sequencing used two gene specific primers and so suffered from PCR bias and over-represented the shorter amplicons. This made quantitative comparisons of alleles of different lengths impossible. We turned to single gene specific primer amplification methods such as UDiTaS and LAM for quantitative analysis because they amplify all outcomes approximately equally. This allowed us to measure the frequency of deletions in *POLB* editing to be 26% of all alleles and only 16% of alleles maintained their wild type allele. 39% of alleles show correct integration of the donor, and the rest would not produce functional protein (structural inversions or deletions). This ability to measure knock-in and knock-out rates concurrently is helpful in understanding the function at the cellular level. In contrast to other studies measuring repair outcomes of a Cas9 DSB (32), we did not detect chromosomal fusions at our break points. However, this may be due to our analysis time point three weeks post-targeting, where alleles could have been selected out of the population. Beyond the utility for quantitative measurements on-target, these single gene primer protocols are powerful for measuring unintended integration of introduced DNA sequences. For example, in treating a mouse model of muscular dystrophy, linear amplification measurements showed the therapeutic AAV unintentionally integrated into the Cas9 break site and throughout the genome (14). Others have recently demonstrated high rates of unintended on and off target integration of AAVs using single primer amplification (38). Replace and Insert donor sequences have the potential to integrate into the target site or off-target into the genome. To our knowledge this is the first work to map and quantify off-target integration or concatenation of donor sequences following NHEJ Insert or Replace targeting. Using a primer on the donor sequence, we detected substantial off-target integration of the donor. Strikingly, none of these off-target integration loci were within 5000 bases of the top 293 predicted Cas9 off-target sites. Rates of off-target integration may be similar for double stranded HDR templates, but to our knowledge off-target integration mapping by linear amplification has not been done after an HDR editing making comparison difficult. Single stranded donor templates are known to integrate off-target less frequently (7), but off-target quantification has mainly relied on integration of large fluorescent cassettes and should be re-evaluated using single primer amplification approaches.

There are currently over 3800 genes known to cause monogenic diseases with mutations often spread across multiple exons (39). Gene editing holds potential to resolve these disorders, however reversing the genetic defects in terminally differentiated or resting cells remains a major challenge (40). HDR is unable to target non-dividing cells (8). NHEJ based Insert targeting and RNA guided transposons (41, 42) are restricted to inserting a sequence at a specific site and cannot remove a mutant region. Engineered RNA-guided recombinases have the ability to delete and insert sequences, but have not yet been tested for replacement of a genomic sequence, and currently have low efficiency (43). Base editor systems (44) can target non-diving cells but currently can only target a subset of disease causing mutation. Prime editing (45) holds potential to be more versatile than base editing, but the effect of delivering a reverse transcriptase to cells needs to be carefully understood across cell types. RNA-targeting CRISPR effectors (46) could target deleterious RNA transcripts but do not solve the underlying genomic issue. Single homology arm donor mediated intron tagging (SATI) (47) has been shown to produce insertions in non-dividing cells in vivo, and could potentially be used for sequence replacements. However this application is yet to be demonstrated and the mechanism and utility across cell types need to be evaluated. Currently, Replace editing is the only technique allowing the efficient exchange of large or small sequences using a repair pathways active in dividing and non-dividing cells. As with any genetic editing that involves a DSB formation, Replace editing should be analyzed by techniques such as linear amplification deep sequencing to monitor and understand the outcomes of repaired alleles.

Taken together this work establishes the Replace method as a viable method for exchanging genomic sequences in human cells. The utilization of NHEJ for the genomic replacements provides a basis for future work to edit slowly dividing and non-dividing cells for gene therapy and in research.

## Materials and Methods

### Data and Methods Availability

Sequencing data is available. Sequence Read Archive (SRA) accession: PRJNA622521. Extended protocols are available: https://www.protocols.io/researchers/eric-danner/publications. Plasmids were submitted to Addgene: https://www.addgene.org/Ralf_Kuehn (#149344-#149354) and a folder of annotated genebank (.gb) files is added as Supplementary File 1. All code used is available on Github: https://github.com/ericdanner. This includes scripts, Jupyter notebooks, and Conda environments.

### DNA Constructs

Cas9-2A-puro targeting plasmid (#488) is Addgene ID 62988 with F1 sequence removed. The AAVS1 targeting fluorescent reporter #208 system was modified from Addgene ID 60431. The neomycinR sequence was modified to a more robust form (48). The RTTA3 gene was replaced by a BFP-pA-Venus-pA where the BFP is flanked by Rosa26 sequences constructed by Gibson Assembly. Guide RNA target sequences were ligated into BbsI cleaved plasmids using synthetic oligonucleotides (Table 1). When more than one guide was necessary the plasmids were combined using Gibson Assembly.

Minicircles are produced in engineered bacteria using arabinose-induced recombination to remove the plasmid backbone (49, 50). The ZYCY10P32T E.Coli strain and the minicircle backbone were purchased from System Bioscience. After cloning in the sequence into the specific minicircle backbone the plasmid is transformed into the ZYCY strain. The 200 ml culture was grown in TB media for 16 hours. Then 200 μl of 20% L-arabinose were added and adjusted to pH 7 and 200 ml LB were added. The culture was then shaken at 32°C for 4 h to induce minicircle formation and slow cell division. An endotoxin free purification kit (Macherey Nagel) was used following the protocol for low copy number plasmids. The resulting product contained plasmid and gDNA contamination. Restriction enzymes cutting the backbone and gDNA were added for 2 hours. Then the resulting fragmented DNA was digested with PlasmidSafe DNase for 16h (Epicure).

### Cell Culture and Targeting

HeLa cells were cultured in DMEM, 10% FBS, 1% Penicillin/Streptomycin and passaged with trypsin every 3-4 days. To generate the fluorescent reporter line plasmid #208 was cloned. Successful integration into the AAVS1 loci generated neomycin resistance. Cells were selected with 0.6 mg/ml G418 for 1 week. Single cells were FACS sorted into a 96 well plate and expanded. Colonies were checked for correct integration by genotyping and a clone with inserts on both alleles was expanded and used. Targeting of Reporter HeLa: 50,000 cells were reverse-transfected with 1.5 μg of Cas9_2A_puro/guide plasmid + 1.5 μg of MC or plasmid complexed with Lipofectamine 3000. The next morning 1.5 μg/ml puromycin was added for 48 hours. Cells were then FACS analyzed. mCherry^+^ cells were single cell sorted into a 96 well plate and expanded for genotyping. For the HDR targeting experiment the guide RNA targeting the Insert site was used together with the donor plasmid.

K-565, a leukemia cell line, were kept in IMDM, 10% FBS, 1% Penicillin/Streptomycin and split every 3 days. For targeting cells were nucleofected using the Lonza 4D strips. 5 x 10^5^ cells were resuspended in nucleofection buffer (51) with 1 μg Cas9/guide plasmid and 3 μg of minicircle and nucleofected using program FF-120. The following day puromycin (4 μg/ml) was added for 48 h.

### Genotyping

For single cell clones or bulk sequencing genomic DNA (gDNA) was extracted by quick extract (Lucigen). PCR amplification was performed with LongAmp Polymerase (NEB) or PrimerStar GXL (Takara). Primer pairs flanking the upstream cut site or downstream cut-site were used. Amplicons were verified by gel extraction and Sanger sequencing. Amplicons from bulk sequencing were cloned into the TOPO vector (Invitrogen) before Sanger sequencing.

Frequency of homozygous and heterozygous integration in HeLa cells was determined by knocking-in mCherry and miRFP670 simultaneously. By measuring mCherry^+^, miRFP670^+^, and double positive cells the homozygous knock-in could be calculated (7).

We used modified ICE analysis for deconvolution of amplicon Sanger trace data derived from unsorted Replace targeted cells (37). The amplicons were made using a primer on the donor sequence and a primer on the genomic sequence flanking the ligated site. The amplicon was cloned into the TOPO vector and individual cloned alleles were Sanger sequenced along with the mixed PCR product. A cloned colony with Replace inserts without any InDels was identified, and this Sanger trace data was used as the ‘wild-type’ reference in ICE analysis.

### Long Read Deep Sequencing and Analysis

https://github.com/ericdanner/REPlacE_Longread

Bulk gDNA of targeted and control cells was amplified by PrimeStar GXL for Polb targeting. The HeLa Deletion Reporter required PCR with OneTaq (NEB) using the high GC content additive to amplify through the very GC-rich CAG sequence. 5 minutes elongation steps were used to reduce PCR bias. Amplicons were cleaned by SPRI beads and quantified by Qubit. The Libraries were pooled and prepared for PacBio sequencing following company protocol. Data Analysis was done using ‘Pipeline Longread’. was done using custom Python scripts for preprocessing and binning of the reads into different structural variants: original exon, replacement, insertion, deletion. Alignments were done with BBmap or MiniMap2 (52) and visualized with IGV (Interactive Genome Viewer). Analysis of alignments was done in R using a modified script from (Github/pigX). Plotting was done in R or Python with a number of the plots included in the Jupyter Notebooks.

### Uni-Directional Targeted Sequencing Sample Preparation

Wild Type and treated cells having had the *POLB* exon 5 targeted showing 50% mCherry expression were used. Samples were prepared either as described in LAM-HTGTS (33) beginning with 500ng of gDNA or based on the Tn5-Uditas protocol (32) beginning with 50ng gDNA. LAM-HTGTS was done generally as published with a few modifications. A single biotinylated gene specific primer was used to amplify 500ng sonicated gDNA (1kb peak) 80x rounds. Streptavidin Dynabeads were found to inhibit PCR so the concentration was reduced to 1/10th and used to capture the amplified sequence. Capture bead-DNA was washed and then the universal primer was ligated on the end. This adapter-ligated sequence was PCR amplified with a universal primer and a nested gene specific primer 30x. We added Nextera adapters by 10x rounds of amplification. Gel extract 300-500bp smear 300-500bp, quantified by Qubit and Bioanalzyer, then sequenced with Illumina MiniSeq. For Tn5 sample preparation we modified the UDiTaS protocol, 50 gDNA was washed 2x with SPRI beads. Tagmentation used hyperactive Tn5 produced by the Max Delbrueck Center protein production facility following published protocols [46]. Samples were tagmented to add the universal primer binding site. Sample was amplified with gene specific primer and universal primer 15x. A nested primer with Illumina adapter sequences was added and followed by PCR 15x. Then Illumina adapters were added with 10x PCR. Amplicons 300-500bp were gel extracted, quantified by Qubit and Bioanalzyer, then sequenced with Illumina MiniSeq.

### Analysis of Uni-Directional Targeted Sequencing

All scripts and notebooks are on github.com/ericdanner/Uditas-Replace. The analysis of the linear amplified sequences was based on the Uditas software. De-multiplexed samples are run through pipeline 1 or pipeline 2. Pipeline 1 generates amplicons of the various expected outputs and does a global alignment using Bowtie2 (53). Reads that align well and cover the ligated junctions are analyzed for InDels. If the samples were prepared with Tn5 they contained UMIs. Unique UMIs are tallied and editing outcomes are quantified. LAM samples do not contain UMIs. In Pipeline 2 the reads are checked for correct on-target priming. The samples are then trimmed using Cutadapt (54) up to the expected break site leaving only the sequence downstream of the break-site. This sequence is aligned globally using Bowtie2 to an index file containing hg38 and the targeting vector.

## Acknowledgments

We would like to thank Julian Clauss for assistance in establishing the Minicircle production. Bora Uyar and Altuna Akalin of the Bioinformatics Core Facility at the Max Delbrueck Center for mentorship. Claudia Quedenau, Tatiana Borodina, and Sasha Sauer of the Max Delbrueck Center Sequencing Core Facility for assistance with PacBio and Illumina sequencing and Hans Peter Rahn from the Max Delbrueck Center FACS core facility for assistance with cell sorting. Julia Huntenburg for crucial feedback on the manuscript. Plasmids <u>AAVS1-SA-2A-NEO-CAG-RTTA3</u> (ID 60431) and pSpCas9(BB)-2A-Puro (PX459) (ID 62988) were gifts from Paul Gadue and Feng Zhang, obtained via Addgene.

## Primer Table

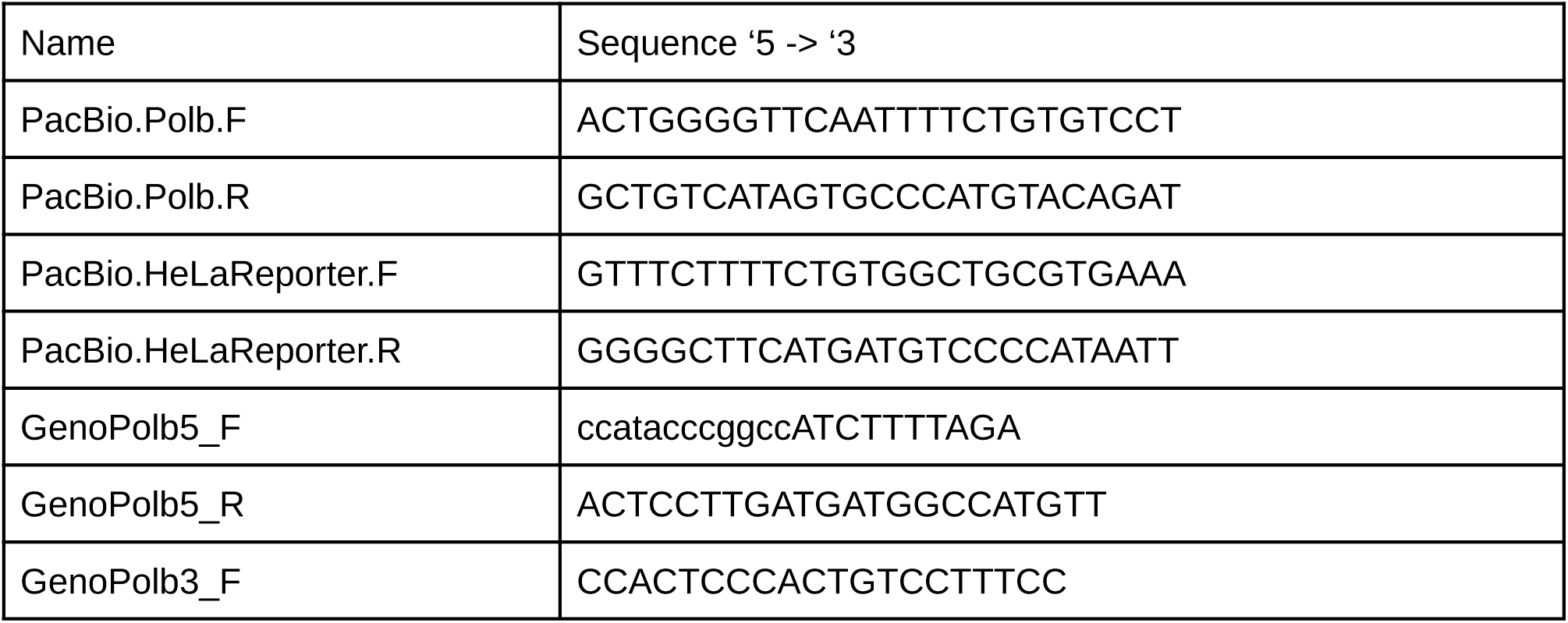
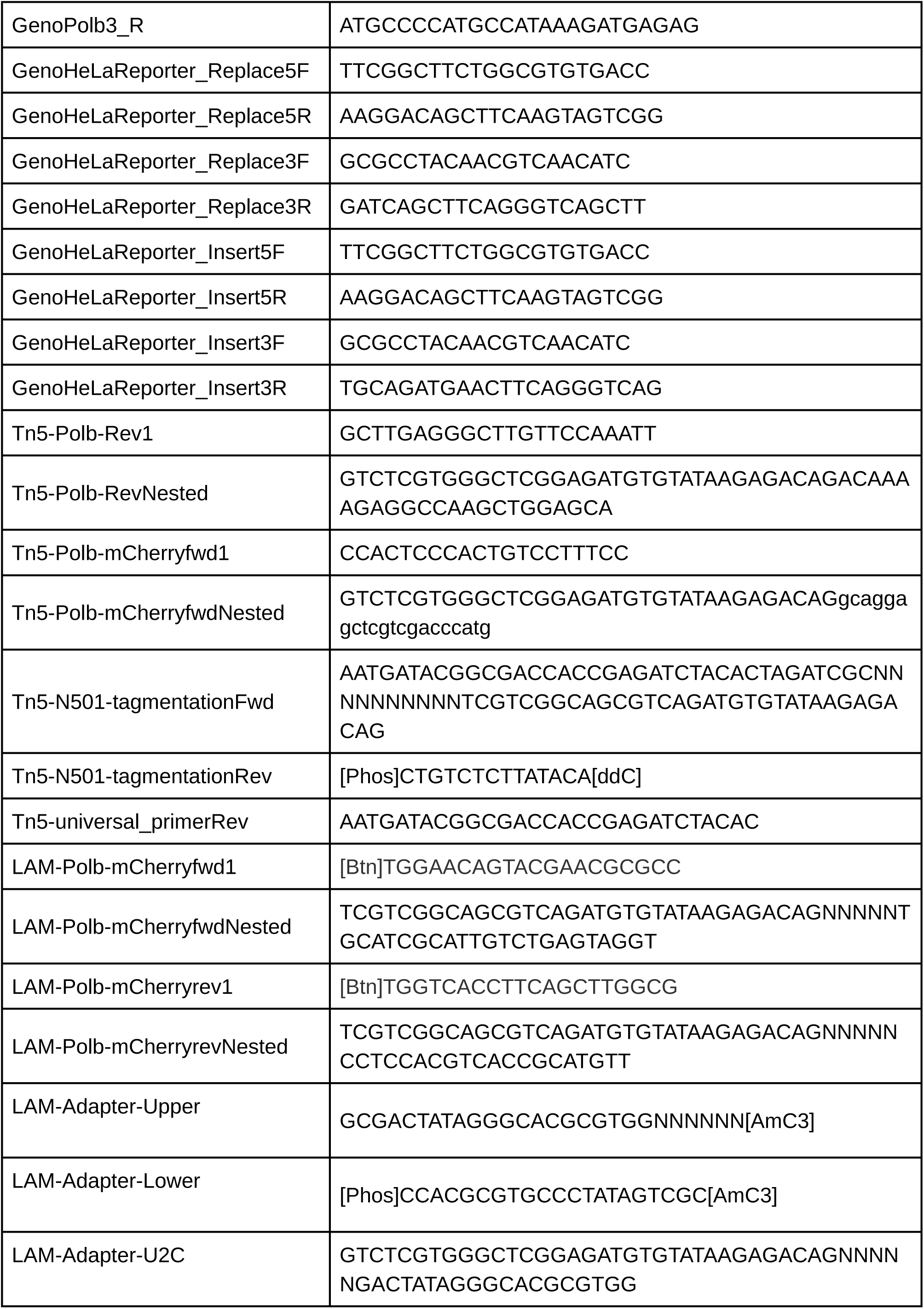

**Supplementary File 1:** .gb sequences zipped for the following annotated files:

MiniCircleProductionPlasmid

Polb_WT

Polb_ReplaceTargeted

Polb_DonorSequenceMC

PolbTargeting_Cas9_guides

HeLaReporter_WT

HeLaReporter_ReplaceTargeted

HeLaReporter_InsertTargeted

HeLaReporter_ReplaceDonorSequenceMC

HeLaReporter_InsertDonorSequenceMC

HeLa_DonorSequencemiRFP670

HeLaReporter_ReplaceCas9_guide LMNA_WT

LMNA_ReplaceTargeted CCNA_WT

CCNA_ReplaceTargeted

## Supplementary Figs

**Supplementary Figure 1:**
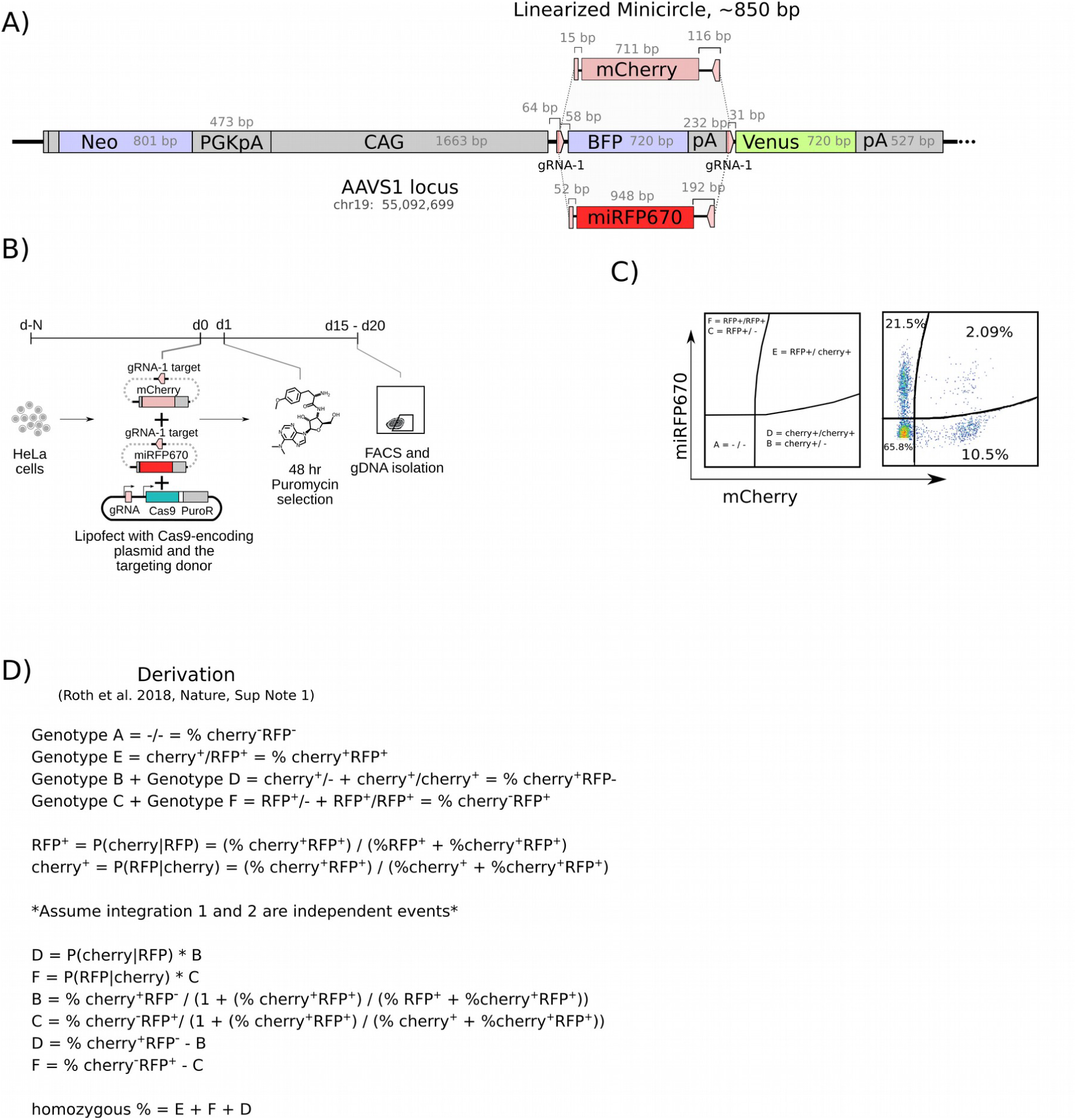
Quantification of homozygous knock-in in HeLa reporter cells. **A.** Quantification strategy uses two otherwise identical constructs with different fluorophores: mCherry and miRFP670 for Replace targeting. **B.** Lipofected reporter cells were selected for 48 hr and analyzed by FACS after 2-3 weeks. **C.** Example of a FACS plot at day 15. **D.** Derivation used to quantify homozygous and heterozygous knock-in events.

**Supplementary Figure 2:**
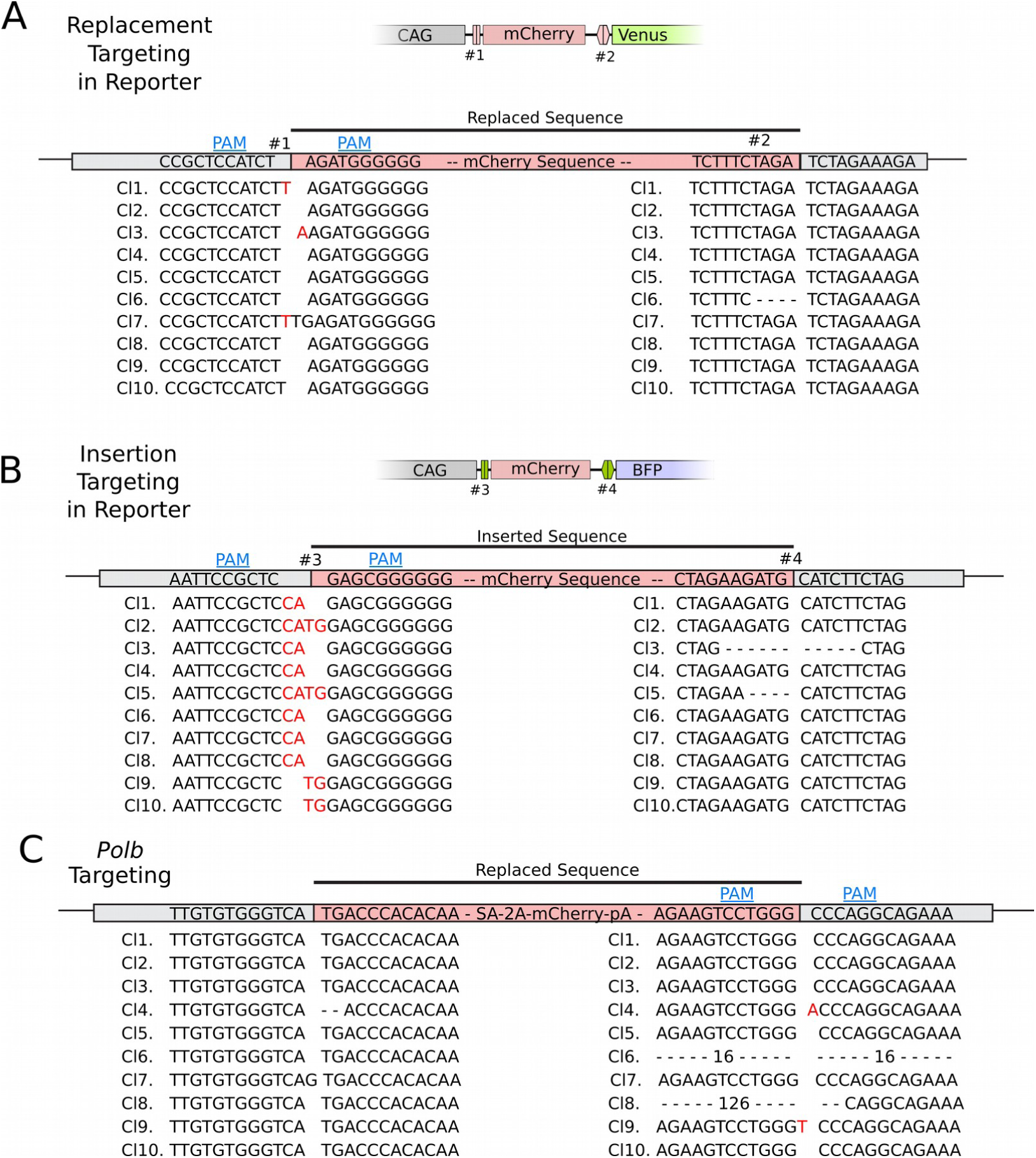
Sanger sequence analysis of targeting experiments shown in Figure 1 and 4. **A.** InDels at the interface of the integrated donor sequence in Replace targeting HeLa reporter clones shown by Sanger sequencing of cloned PCR products. **B.** InDels at the interface of the integrated donor sequence in Insert targeting HeLa reporter clones shown by Sanger sequencing of cloned PCR products. **C.** InDels at the interface of the integrated donor sequence in replace targeting *POLB* exon 5 in K562 clones shown by Sanger sequencing of cloned PCR products.

**Supplementary Figure 3:**
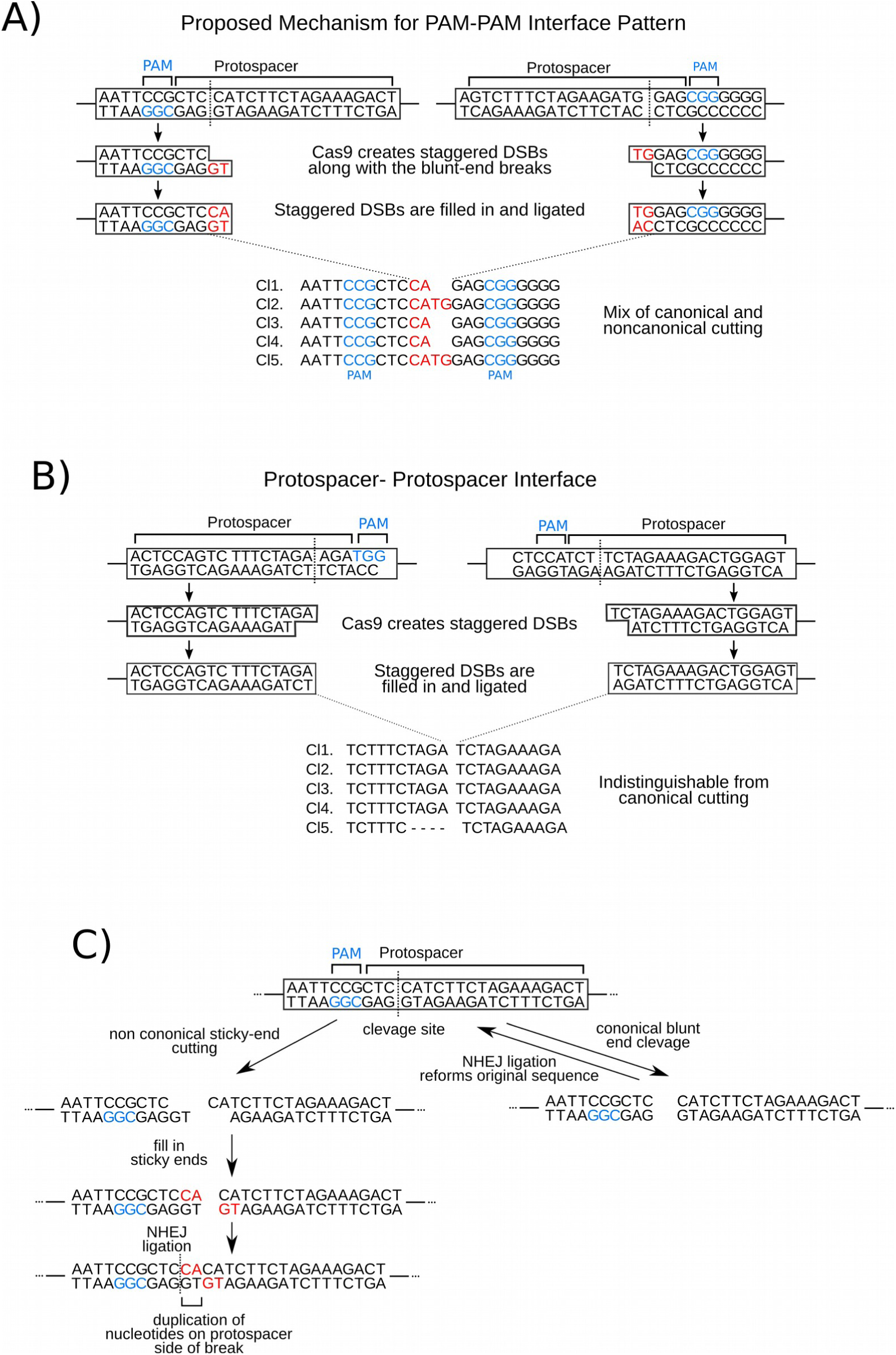
Effects of staggered cuts on Replace targeting. **A.** For Replace or Insert targeting upon the ligation of two PAM-PAM ends the product will show insertions when formed by a staggered cut. **B.** The protospacer-protopacer junction will not show the effects from a staggered cut even when the staggered cut occurs frequently. **C.** Staggered cuts at a single target site produce the insert and halt further DSB formation.

**Supplementary Figure 4:**
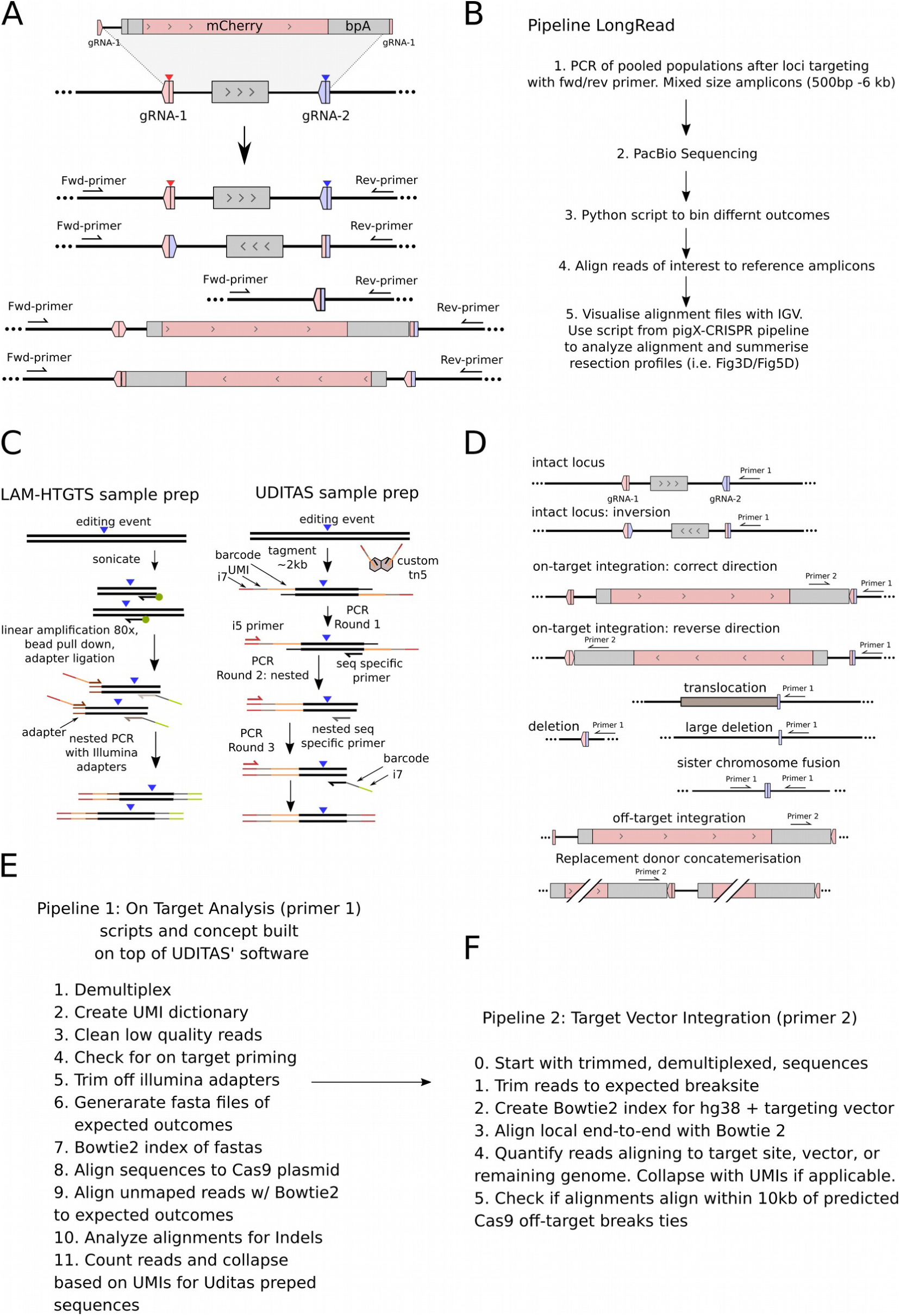
Deep sequencing algorithms. **A.** Two primer amplification captures some structural variants produced by Replace targeting. These products are then used for long read sequencing. **B.** Pipeline Long read overview. **C.** Linear amplification as prepared using the LAM-HTGTS or UDITAS protocols. **D.** Complex outcomes that are captured by linear amplification preparations using a locus-specific primer (primer 1) or replace donor sequence primer (primer 2). **E.** Pipeline 1 overview for on target loci analysis. **F.** Pipeline 2 overview showing donor sequence integration analysis.

**Supplementary Figure 5:**
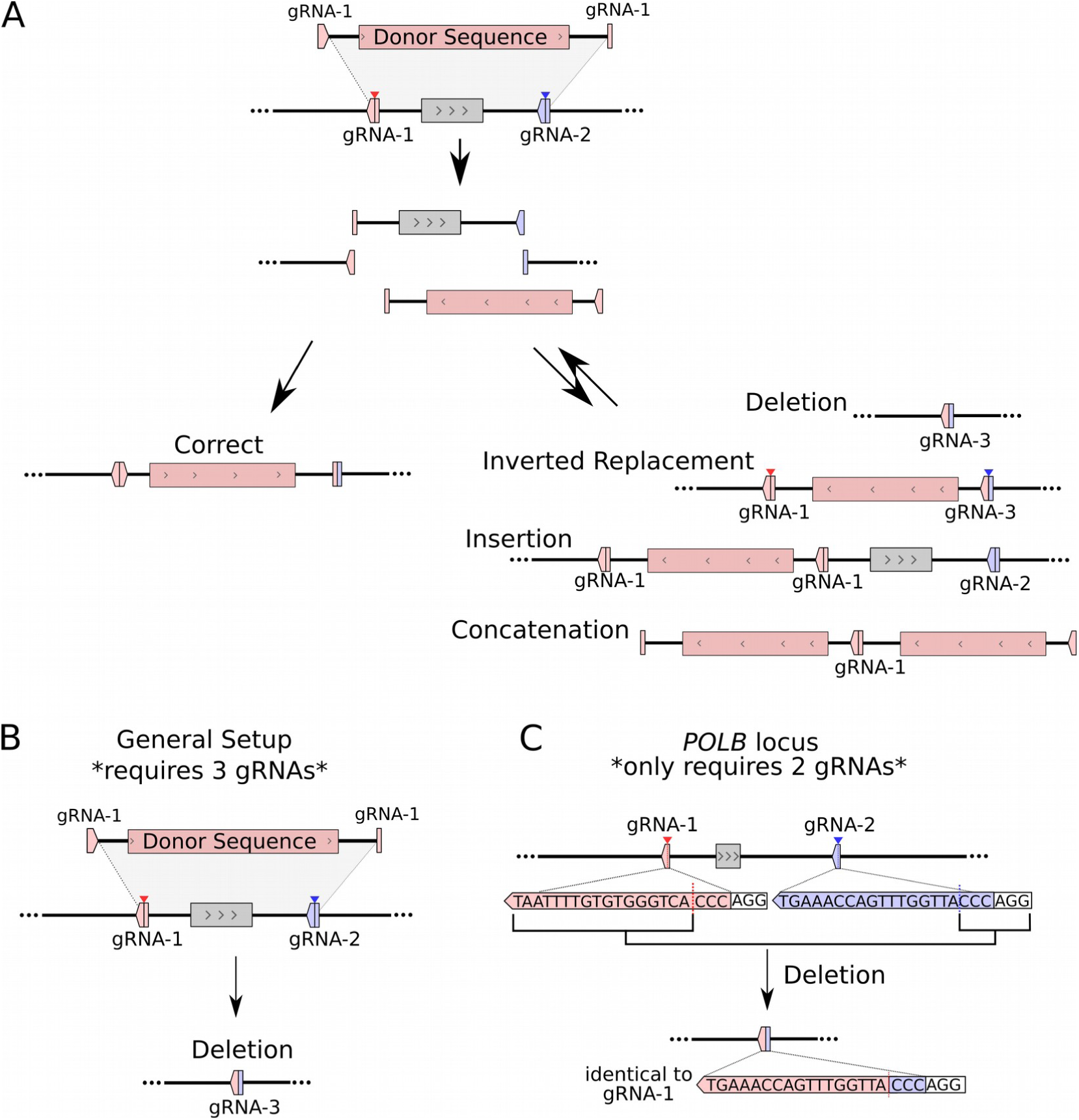
Designed directionality in Replace Targeting. **A.** Using the principles from Insert targeting by Suzuki et al (2016) we designed Replace donor directionality with using any homology sequences. The donor sequence is linearized or excised by gRNA-1 and the genome cut by gRNA-1 and gRNA-2. The gRNA-1 site on the donor is oriented so that if the donor sequence is inserted in an undesired way it reforms a cut site. **B.** The general setup requires two guides to cut the genome and gRNA-3 to cut the site formed by a deletion. **C.** For *POLB, CCNA1, LMNA Replace targeting* we only used gRNA-1 and gRNA-2, and avoided the need of gRNA-3. We selected gRNAs in which the first 3 nucleotides of the protospacer are identical for both gRNAs, so the deletion reforms gRNA-1’s recognition sequence.

**Supplementary Figure 6:**
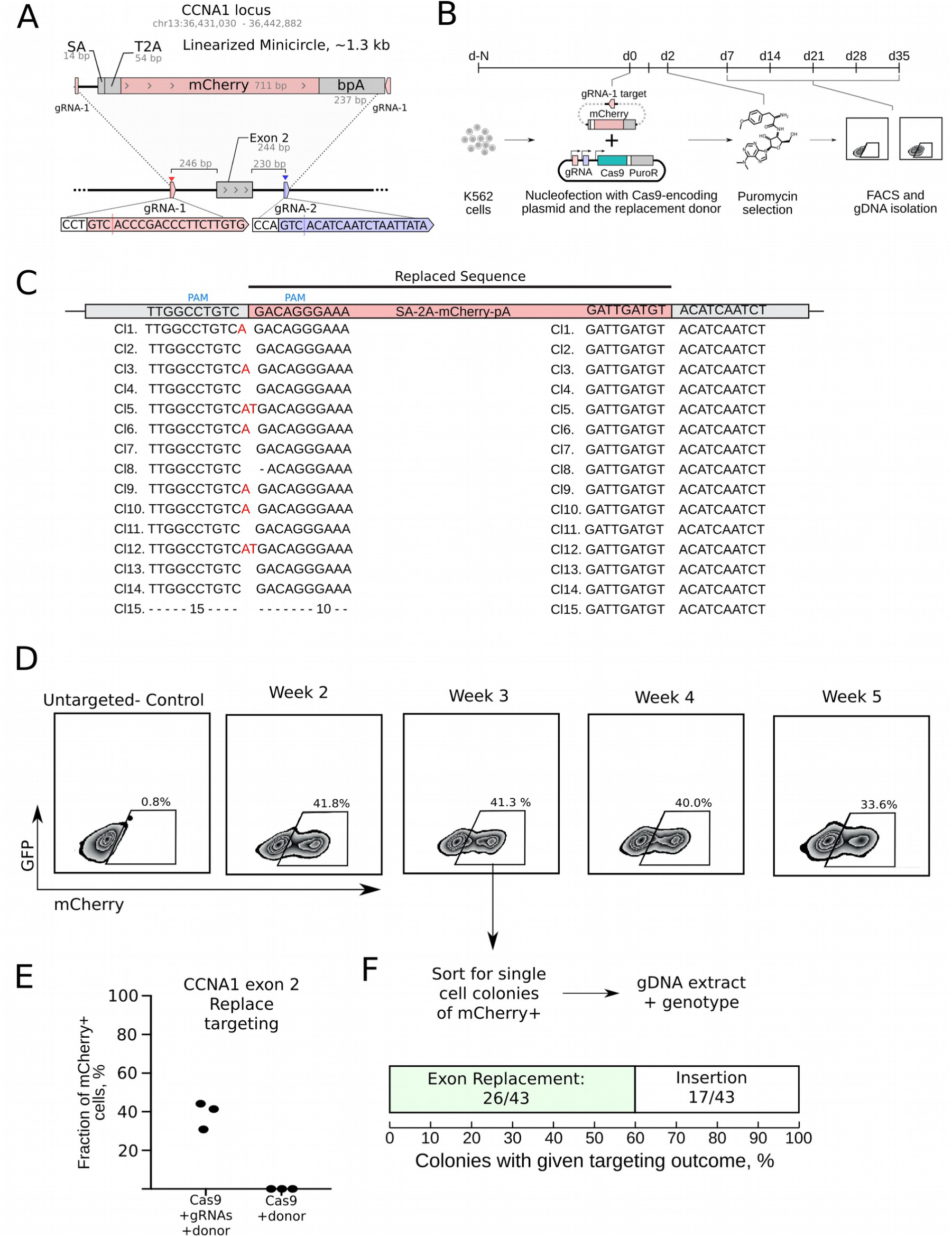
Replace targeting of *CCNA1* exon 2 in K562 cells. **A.** Targeting design for Replacing exon 2 with a mCherry reporter including a splice acceptor (SA), T2A self-cleaving peptide and a polyadenylation site (bpA) using a minicircle. **B.** Nucleofected K562 cells were selected with puromycin for 48 hours and then analyzed weekly by FACS. **C.** InDels at the interface of the integrated donor sequence in Replace targeted cells shown by Sanger sequencing of cloned PCR products. PAM-PAM junctions show familiar insertion patterns (red) caused by non-canonical cutting. **D.** Stable expression of mCherry in transfected cells over time. **E.** Efficiency of Replace targeting at *CCNA1* exon 2. (Duplicate of Figure 4G) **F.** Genotyping results for mCherry^+^ single cell derived colonies show replacement occurred in 60% of mCherry^+^ cells.

**Supplementary Figure 7:**
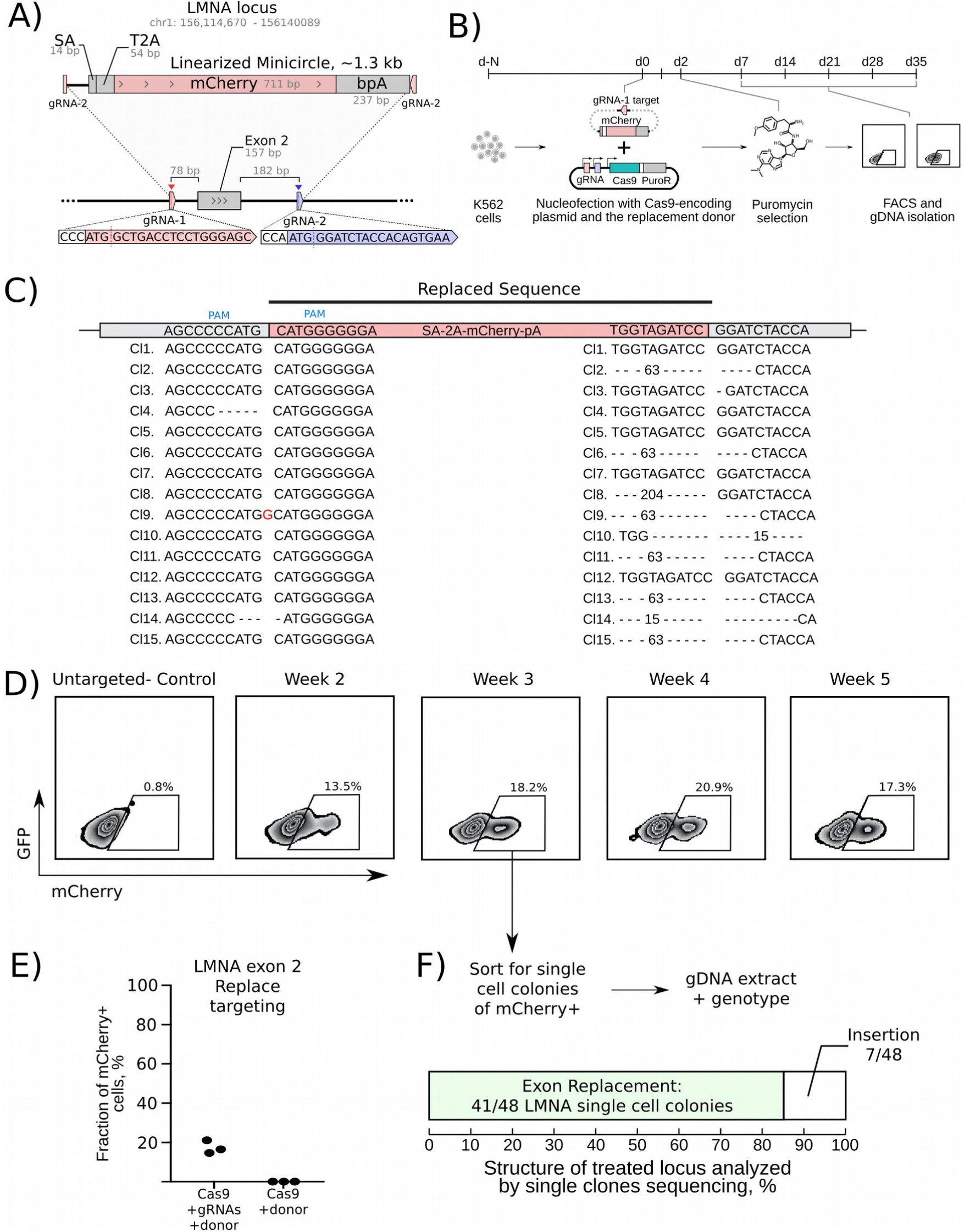
Replace targeting of *LMNA* exon 2 in K562 cells. **A.** Targeting design for Replacing exon 2 with a mCherry reporter including a splice acceptor (SA), T2A self-cleaving peptide and a polyadenylation site (bpA) using minicircle donors. **B.** Nucleofected K562 cells were selected with puromycin for 48 hours and then analyzed weekly by FACS. **C.** InDels at the interface of the integrated donor sequence in Replace targeted cells shown by Sanger sequencing of cloned PCR products. **D.** Stable expression of mCherry in transfected cells over time. **E.** Efficiency of Replace targeting at *LMNA* exon 2 (Duplicate of Figure 4H). **F.** Genotyping results for mCherry^+^ single cell derived colonies show replacement occurred in 85% of mCherry^+^ cells.

**Supplementary Figure 8:**
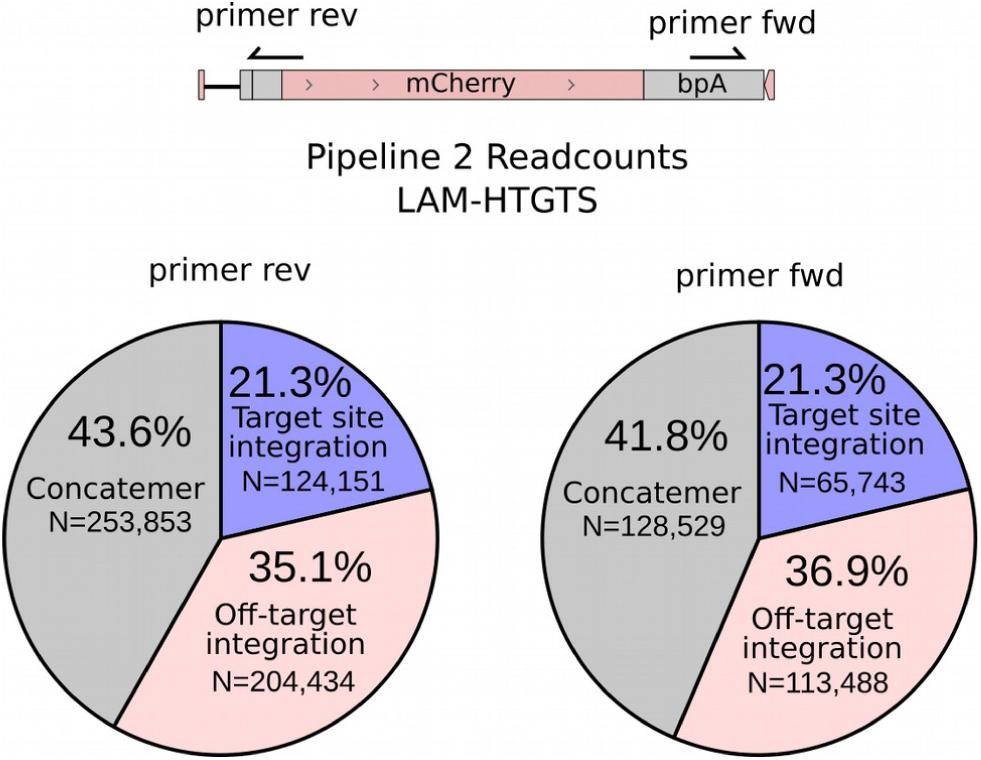
LAM off target integration for Replace targeting of *POLB* exon 5 in K562 cells.

